# A genetic-based algorithm for recovery: A pilot study

**DOI:** 10.1101/166181

**Authors:** Craig Pickering, John Kiely, Bruce Suraci, Charlie Quigley, Jake Watson

## Abstract

Exercise training creates a number of physical challenges to the body, the overcoming of which drives exercise adaptation. The balance between sufficient stress and recovery is a crucial, but often under-explored, area within exercise training. Genetic variation can also predispose some individuals to a greater need for recovery after exercise. In this pilot study, 18 male soccer players underwent a repeated sprint training session. Countermovement jump (CMJ) heights were recorded immediately pre-and post-training, and at 24-and 48-hours post-training. The reduction in CMJ height was greatest at all post-training time points in subjects with a larger number of gene variants associated with a reduced exercise recovery. This suggests that knowledge of genetic information can be important in individualizing recovery timings and modalities in athletes following training.

## Introduction

Exercise training produces a variety of acute physiological challenges to the body, perturbing homeostasis and inducing a stress response, the overcoming of which leads to exercise adaptation. Successful adaptation is comprised of the accumulation of periods of stress-induced response (training), and periods of recovery, which takes place away from exercise (Bishop et al., 2008). This relationship is finely balanced, and if there is insufficient time between training sessions for recovery to occur, the athlete increases their risk of undue accumulation of fatigue, potentially resulting in acute underperformance, injury (Mair et al., 1996), illness (Schwellnus et al., 2016), and eventually non-functional overreaching and unexplained underperformance syndrome (UPS) (Soligard et al., 2016). These indicators of maladaptation are common in athletes, with 10-20% of endurance athletes suffering from UPS each year (Budgett, 2000). Overuse injury incidence is also frequent, with rates between 37% and 85% reported, depending on the sport (DiFiori et al., 2013; Wilber et al., 1995). The prevalence of these symptoms of stress-recovery imbalance at epidemic proportions indicates that there is a mismatch in our knowledge of how to create stress and how to recover from it. Indeed, a search on PubMed yields over 37,000 papers with “exercise training” in the title and abstract. A similar result with “exercise recovery” as the search field only results in 13,000 papers.

At the cellular level, the physiological challenges induced by exercise include increased oxidative metabolism within the mitochondria, leading to increased formation of reactive oxygen species (ROS) (Fisher-Wellman & Bloomer, 2009). Under normal, non-exercise conditions, the body can neutralise ROS through its endogenous antioxidant defence system, which is comprised of enzymes such as superoxide dismutase (Belviranli & Gokbel, 2006). However, when ROS production is increased through exercise, an imbalance between ROS production and neutralisation occurs, leading to elevated oxidative stress (Fisher-Wellman & Bloomer, 2009). In turn, this elevates lipid peroxidation and tissue damage. Mechanical load also increases muscle damage (Baumert et al., 2016), initiating an inflammatory response partially driven by pro-inflammatory cytokines such as interleukin-6 (IL-6) and tumour necrosis factor (TNF) (Yamin et al., 2008).

These changes at the molecular and cellular level drive the whole-body symptoms of under-recovery that athletes and coaches are well aware of. Increases in plasma IL-6 levels occur following exercise (Robson-Ansley et al., 2007), and administration of exogenous IL-6 into athletes induces feelings of fatigue and impairs performance (Robson-Ansley et al., 2004). Increased IL-6 is also a risk factor for the development of UPS (Robson, 2003). Both TNF and IL-6 can act on the central nervous system (Ament & Verkerke, 2009), potentially decreasing the drive to exercise. Increased ROS and oxidative stress are also associated with a decrease in physical performance (Powers & Jackson, 2007), and increased feelings of muscle soreness following exercise (Konig et al., 2001).

Recently, a number of best practices for recovery have been put forward (Soligard et al., 2016; Schwellnus et al., 2016, Leeder et al., 2011, Bishop et al., 2008). However, there is likely considerable inter-individual variation in the time course of exercise recovery (Nosaka et al., 1996). This is partially governed by genetic variation between individuals, with several single nucleotide polymorphisms (SNPs) already identified as potentially impacting the speed of post-exercise recovery (Baumert et al., 2016). Knowledge of this variation may enable athletes and support staff to create individualised recovery interventions based on the identification of individuals at increased risk of exercise induced muscle damage or oxidative stress (Del Coso et al., 2017).

Whilst this emerging research of the impact of genotype on exercise recovery is interesting, the translation of these findings to the field are currently under-explored. Given the proposed impact of genetic polymorphisms on exercise recovery, the purpose of this study was to attempt to bridge this gap, by determining whether a seven SNP algorithm successfully differentiated between the recovery speed of male soccer players. It is believed that individuals possessing a greater number of alleles associated with increased oxidative stress, muscle damage or inflammation would see a greater reduction in neuromuscular function post-training, and that this reduction would take longer to abate relative to those individuals in possession of a more favourable genetic profile.

## Methods

### Subjects

18 male soccer players aged between 16-19 years of age from a college soccer academy volunteered to participate in this study. Each player had an average of 11 years’ football training experience, and was actively competing in the English College Football Association Premier League.

### Methodology

As part of their normal soccer training, and following 24-hours rest, the subjects took part in a repeated sprints testing session. Both before, and immediately upon completion of this session, the subjects underwent Countermovement Jump (CMJ) testing. This test was repeated at 24-and 48-hours post-training to monitor their recovery status. Subjects were familiarised to all tests as they are regularly used during their normal soccer training. Within the CMJ trials, subjects undertook three trials at each time point, which were averaged to give a mean score. Subjects had two minutes’ recovery between trials. Prior to the initial exercise bout and subsequent testing, players carried out a standardised 15-minute warm up, consisting of pre-activation exercises and dynamic drills. The initial exercise bout was comprised of two sets of seven 25m sprints undertaken outdoors on a soccer pitch. The recovery period was 30 seconds between sprint reps, and 5 minutes between sets.

The CMJ was chosen as it is a reliable and valid measure of neuromuscular fatigue (Cormack et al., 2008a, 2008b, 2008c; McLean et al., 2010; Gathercole et al., 2015) that is widely used in sporting settings (Taylor et al., 2012). The CMJ was measured using Optojump (Microgate, Italy), a valid and reliable tool for the assessment of vertical jump height in the field (Glatthorn et al., 2011). Prior to undertaking each CMJ, subjects were instructed to keep their hands on their hips throughout the jump to eliminate any influence of arm swing. If the arms lost contact with the hips, the jump was classed as a no-jump, and an additional jump was performed following two-minutes recovery. In each CMJ trial, subjects began standing upright, then performed a fast downwards eccentric action followed immediately by a jump for maximal height. Individual results were expressed as height jumped in centimetres (cm).

### Genetic Testing

Alongside the training programme, subjects underwent genetic testing by DNAFit Ltd; this occurred via a sterile buccal swab. The samples were sent to IDna Genetics Laboratory (Norwich, UK), where DNA was extracted and purified using the Isohelix Buccalyse DNA extraction kit BEK-50 (Kent, UK), and amplified through PCR on an ABI7900 real-time thermocycler (Applied Biosystem, Waltham, USA). Through this process, genetic information regarding SNPs believed to affect post-exercise recovery speed (*CRP* rs1205, *GSTM1* & *GSTT1* INDEL, *IL-6* -G174C rs1800795, *IL-6R* rs2228145, *SOD2* rs4880, *TNF* G-308A rs1800629) was determined. The DNAFit test uses an algorithm to stratify subjects into “slow”, “medium” or “fast” recovery speed by utilising a Total Genotype Score (TGS) method. Each allele is given a score of between 0 and 4 points depending on the expected magnitude of its impact on post-exercise recovery speed. The strength of the rating was based on the evidence from cumulative literature results averaged over time. The sum of these points was combined to give an overall score. This method is identical to Jones et al. (2016), and similar to the methods used in other studies utilising genetic algorithms (Ruiz et al., 2009; Meckel, Ben-Zaken, Nemet, Dror & Eliakim, 2014). An overall score of 40% or less is classed as a fast genetic recovery speed. Scores of 41-60% are classed as a medium genetic recovery speed. A score of >60% is classed as a slow genetic recovery speed. The athletes were blinded to their genetic results until completion of the final testing.

### Statistical Analysis

As this is not a genetic association study, but an observational study into the impact of TGS on exercise recovery, gene-by-gene analysis was not carried out. Instead, we focused on data pertaining to exercise recovery, and compared this to TGS group. Means and standard deviations were calculated for whole group and sub-groups for both pre-and post-training (0h, 24h and 48h) test scores. CMJ height at the three post-training time points was converted to a percentage of pre-training height. Given the small sample size, and the fact that significance testing is both sensitive to low sample sizes, and doesn’t inform as to the magnitude (Buccheit 2016), we instead calculated effect sizes (Cohen’s d) between the groups at the three post-training time points. The thresholds used were 0.2 (trivial), 0.5 (small), 0.8 (moderate), >0.8 (large) (Cohen 1988). Data were analysed using Microsoft Excel 15.29 (Microsoft Corporation, Redmond, WA, USA). All data are reported as mean ± SD.

## Results

Overall, 12 subjects were classed as having a fast recovery speed, with 6 subjects having a medium recovery speed. No subjects were found to have a slow recovery speed; based on an analysis of 17,000 samples tested by DNAFit Ltd (Pickering, unpublished data), approximately 6% of all individuals within a population would be expected to be in the slow group. Within this sample population, we would expect one subject to be in this group; the lack of this TGS is therefore not unusual. Table 1 shows the absolute CMJ results for the fast and medium genetic recovery speed groups. Figure 1 illustrates the between group differences as a percentage over the 48-hour period following the exercise bout, along with effect sizes.

**Table 1.**
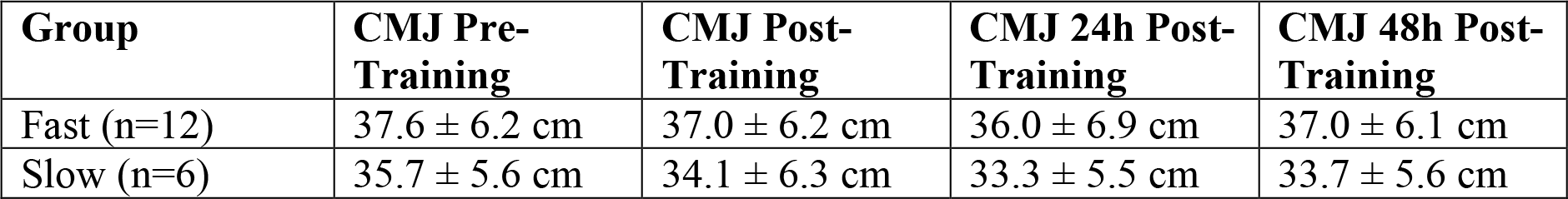
CMJ values for both groups across all time points.

| Group | CMJ Pre-Training | CMJ Post-Training | CMJ 24h Post-Training | CMJ 48h Post-Training |
| --- | --- | --- | --- | --- |
| Fast (n=12) | 37.6 ± 6.2 cm | 37.0 ± 6.2 cm | 36.0 ± 6.9 cm | 37.0 ± 6.1 cm |
| Slow (n=6) | 35.7 ± 5.6 cm | 34.1 ± 6.3 cm | 33.3 ± 5.5 cm | 33.7 ± 5.6 cm |

**Figure 1.**
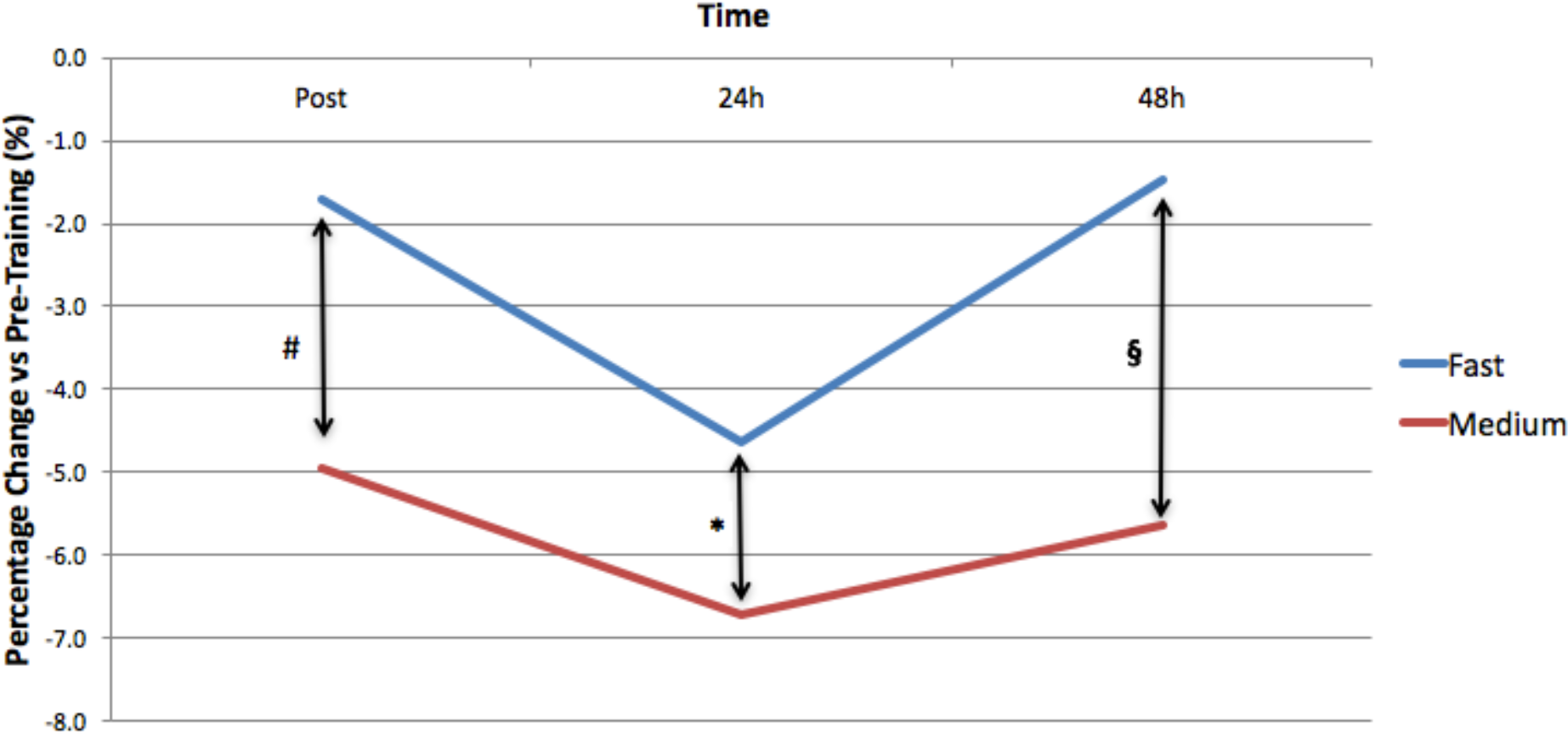
Percentage change in CMJ height immediately post-training, 24h post-training, and 48h post-training as a percentage relative to pre-training values. Effect sizes are # = 0.7 (medium), * = 0.5 (medium), § = 1.0 (large).

## Discussion

The results of this study indicate that rate of recovery, as measured by CMJ, is modified by a total genotype score comprised of seven SNPs thought to impact exercise recovery. Overall, players in the “fast” genetic recovery speed group saw a smaller reduction in CMJ height relative to those players in the “medium” genetic recovery speed group, and were closer to baseline score following 48-hours of recovery. Immediately upon completion of the exercise bout, and 24 hours later, the magnitude of this effect was medium. Forty-eight hours after training, this effect had grown in magnitude to large. This suggests that these seven SNPs impact recovery speed, such that individuals with more favourable alleles suffer a smaller percentage loss in CMJ height after repeated sprints, and have regained a greater percentage of their pre-training CMJ height 48 hours post-training.

The ability to predict the recovery time needed following intense exercise may be useful for several reasons. Firstly, it ensures that optimal recovery time can be given to individual athletes, reducing the fatigue that will accumulate across a training programme. This will ensure that the athlete is not placed at undue risk of suffering from injuries, of which the risk increases under fatigue, and may guard against the developing of unexplained underperformance syndrome. It may also be useful when planning the final physical conditioning session before a competition, with players genetically predisposed to slower recovery speeds having a longer rest period pre-competition. Finally, it may increase the motivation of players to carry out the correct recovery modalities post-training or post-competition, particularly if they are shown to have a slower recovery speed. However, the results from nutrigenetic research indicate that, at present, individuals rarely make behavioural changes based on genetic information (McBride et al., 2010); whether this is the case in highly motivated sports people is unclear.

The SNPs that comprise the genetic algorithm used appear to influence both the inflammatory response to exercise and the ability to tolerate oxidative stress. *IL-6* -G174C (rs1800795) has been shown to influence creatine kinase (CK) levels following eccentric exercise, with the C allele of *IL-6* associated with higher levels (Yamin et al., 2008; Lappalainen 2009). The *IL-*6 C allele is also associated with increased post-exercise plasma IL-6 (Huuskonen et al., 2009), which is associated with increased fatigue in athletes (Robson-Ansley et al., 2007). *IL-6R* (rs2228145) is also associated with post-exercise IL-6 levels, with the C allele associated with higher concentrations (Reich et al., 2007). Increases in plasma IL-6R concentrations are associated with increased C-Reactive Protein (CRP) levels post-exercise, as well as increased feelings of fatigue in athletes (Robson-Ansley et al., 2009). The *TNF* G-308A (rs1800629) polymorphism is associated with plasma CRP levels following aerobic exercise, with AA genotypes having higher CRP levels (Lakka et al., 2006). *CRP* (rs1205) alters plasma CRP concentrations, a common exercise recovery marker (Miles et al., 2007; Ingram et al., 2009), with G allele carriers having significantly higher CRP levels than AA genotypes (Eiriksdottir et al., 2009).

*SOD2* (rs4880) encodes for manganese superoxide dismutase (MnSOD), which supports the dismutation of mitochondrial superoxide radicals into hydrogen peroxide and oxygen (Li et al., 2005). The T allele of this SNP is associated with increased CK post-exercise (Akimoto et al., 2010; Ahmetov et al., 2014), although this relationship is complex and potentially modified by a subject’s habitual antioxidant nutrient intake (Li et al., 2005). *GSTM1* (glutathione S-transferase M1) and *GSTT1* (glutathione S-transferase T1) are insertion/deletion polymorphisms, with deleted genotypes having poor activity of the enzymes. Whilst these polymorphisms are well studied with regards to health and diet interactions (Palli et al., 2004), only one study has examined their impact on post-exercise muscle damage markers, with no significant differences between the inserted or deleted genotypes (Akimoto et al., 2010). These SNPs were included in the algorithm based on their theoretical impact on exercise recovery (Evelo et al., 1992; Vani et al., 1990).

The identification of athletes who may be genetically predisposed to increased recovery times can lead to the use of targeted recovery modalities. These include nutritional interventions. Phillips et al. (2003) reported that 14 days of supplementation with vitamin E, omega-3 and flavonoids blunted the release of IL-6 and CRP following eccentric exercise. Similar results have been reported from other studies (Satoshi et al., 1989; Jouris et al., 2011; Sacheck et al., 2003). Other interventions that may enhance recovery include cold water immersion (CWI) and the use of compression garments, although the results are currently equivocal (Leeder et al., 2011; Bleakley & Davison, 2009; Ascensao et al., 2010; Jakeman et al., 2009; Duffield et al., 2010; Duffield et al., 2008; Jakesment et al., 2010). It should be noted that exercise adaptation relies on the application of stress to the body, and the use of antioxidant supplements and CWI may blunt this adaptation (Draeger et al., 2014; Yamane et al., 2015), or in some cases reduce speed of recovery (Close et al., 2006).

There are some potential limitations within this study. The population used is very modest (n=18). A standard soccer team is often comprised of 20-25 players; although at any given time some players may be injured and unable to train. The sample size in this study therefore represents a realistic size that coaches and practitioners may encounter in the real world. It is also representative of sample sizes often used in genetic pilot studies (Loy et al., 2015). However, we recognise that this research requires replication in a larger cohort, which we intend to carry out. In addition, this pilot study had no subjects with a TGS that suggested they had a slow genetic recovery speed, only medium and fast. The result of having a slow recovery speed is uncommon, with approximately only one subject for every sixteen expected to be in this category. This requires very large sample sizes in order to recruit sufficient subjects into this category. The subjects were all male, so it in unclear whether the results would be the same in females. Finally, it must be recognised that the use of this algorithm represents a crude measure, as many other SNPs doubtless impact exercise recovery. However, the results of this pilot study are both novel and promising, such that further research in this field should enable the development of personalised recovery guidelines.

### Conclusion

It can be seen that there is considerable variation in the rate of recovery from a repeated sprint exercise in a group of well-trained male soccer players. The use of a seven SNP genetic algorithm aids in the identification of those players who may require longer recovery times between intense exercise bouts, or who may most benefit from targeted recovery interventions. These findings replicate those of Del Coso et al. (2017), although the SNPs utilised vary. The implications of these findings suggest that knowledge of genetic information may be important in individualising recovery timings and modalities in athletes. Future research is required to replicate these findings in a larger cohort, as well as in females, but nevertheless the results potentially herald a further step towards an individualised training process, potentially revolutionising the field of exercise recovery.

